# Switching the left and the right hearts: A novel bi-ventricle mechanical support strategy with spared native single-ventricle

**DOI:** 10.1101/2022.12.12.519951

**Authors:** Emrah Şişli, Canberk Yıldırım, İbrahim Başar Aka, Osman Nuri Tuncer, Yüksel Atay, Mustafa Özbaran, Kerem Pekkan

## Abstract

Mechanical circulatory support (MCS) is used as a bridge-to-heart transplantation for end-stage failing Fontan patients with single-ventricle (SV) circulation. Donor shortage and complexity of the single-ventricle circulation physiology demands novel circulatory support systems and alternative solutions. An out-of-the-box circulation concept in which the left and right ventricles are switched with each other inspired a novel bi-ventricle MCS configuration for the “failing” Fontan patients. In the proposed configuration, the systemic circulation is maintained by a conventional mechanical ventricle assist device while the venous circulation is delegated to the native SV. This approach spares the SV and puts it to a new use at the right-side providing the most needed venous flow pulsatility. To analyze its feasibility and performance, 8 realistic Fontan circulation scenarios have been studied via a multi-compartmental lumped parameter cardiovascular model (LPM). Model is developed specifically for simulating the SV circulation and validated against pulsatile mock-up flow loop measurements for the ideal (*Fontan*), failed (*VD*) and assisted Fontan (*PVR-cmcs*) scenarios. The proposed surgical configuration maintained the cardiac index (3-3.5 l/min/m^2^) providing a normal mean systemic arterial pressure. For a failed SV with low ejection fraction (EF=26%), representing a typical systemic failure, proposed configuration introduced a venous/pulmonary pulsatility of ∼28 mmHg and a drop of 2 mmHg in central venous pressure (CVP) with acceptable pulmonary artery pressures (17.5 mmHg). In the pulmonary vascular resistance (PVR) failure model, it provided approximately 5 mmHg drop in CVP with venous/pulmonary pulsatility reaching ∼22 mmHg. For high PVR failure case with a healthy SV (EF = 44%) pulmonary hypertension is likely to occur, indicating a need for precise functional assessment of the failed-ventricle before it is considered for the proposed arrangement. Comprehensive *in vitro* and *in silico* results encourage this concept as an economical alternative to the conventional bi-ventricle MCS pending animal experiments.

## 1. Introduction

Each year, about 8 in a thousand babies are born with a clinically significant congenital heart defect^1,2^. Single-ventricle (SV) heart defects are among the most serious congenital complications requiring a series of very complex palliative surgical reconstructions with the aim to achieve an optimally working single-ventricle circulation to compensate the missing right-heart. The third surgical stage of this series is the Fontan procedure^3^ first performed in 1971. Following this pioneering surgical procedure, advances in pediatric cardiac surgery have resulted in reduced morbidity and mortality in this vulnerable patient group^4–8^. Unfortunately, the current surgical therapy is palliative; as the child grows, due to vascular remodeling and hemodynamic adaptations, this complex and surgically reconstructed physiology gradually fails, finally leading to severe heart failure at late adulthood. Over the years the number of adult Fontan survivors waiting for heart transplantation have increased dramatically, with severe complications related to the gastrointestinal system, including feeding disorders, liver dysfunction, protein losing enteropathy and plastic bronchitis^9^.

Steadily increasing number of “adult Fontan survivors” with poor quality of life is a major health problem^10,11^, as almost all patients eventually require either heart transplantation or mechanical circulatory support (MCS), during late adulthood^12^. Shortage of donor organ supply has made MCS an inevitable surgical tool for bridge-to-transplantation to improve a patient’s transplant candidacy^13,14^. While a variety of surgical concepts using pulsatile-or continuous flow MCS devices are being proposed^15–19^, novel approaches and breakthrough devices are still desired to address the physiology-related limitations of the single-ventricle circulation. For example, our group recently proposed an implantable Fontan ventricle assist device without external power and without inlet/outlet tubing^20^. For the right-heart MCS, compared to the left side, the low pulmonary vascular resistance (PVR) requires a high-volume/low pressure pump^21^. This requirement is not necessarily compatible with the existing conventional MCS devices that can supply higher afterloads^22^. Most importantly, the clinically available MCS systems could not achieve physiological pulsatility levels at the right side, which is essential to preserve the venous, pulmonary, and lymphatic function of Fontan patients^11^. Therefore, in this study, we introduce a novel Fontan-MCS modification that can provide the desired physiological “native right ventricle-like” pulsatile flow to support and gradually heal the Fontan failure. Hemodynamic characteristics of this concept was analyzed focusing primarily on the patients with their early stage of failed Fontan circulation that display systemic heart failure (New York Class II and III), in whom a left ventricular mechanical assist device is normally considered as a bridge-to-heart transplantation (Figure 1a). The concept was also tested for the high-PVR Fontan failure model. We hypothesize that in combination with the conventional left ventricle assist device therapy, rather than discarding the native SV, sparing it for the right heart support would result the desired venous hemodynamics (Figure 1b). Furthermore, in some patients, where a bi-ventricle MCS support is essential, this approach will provide significant cost savings that is much significant for resource-limited settings and third-world countries with limited access to MCS devices.

**Figure 1.**
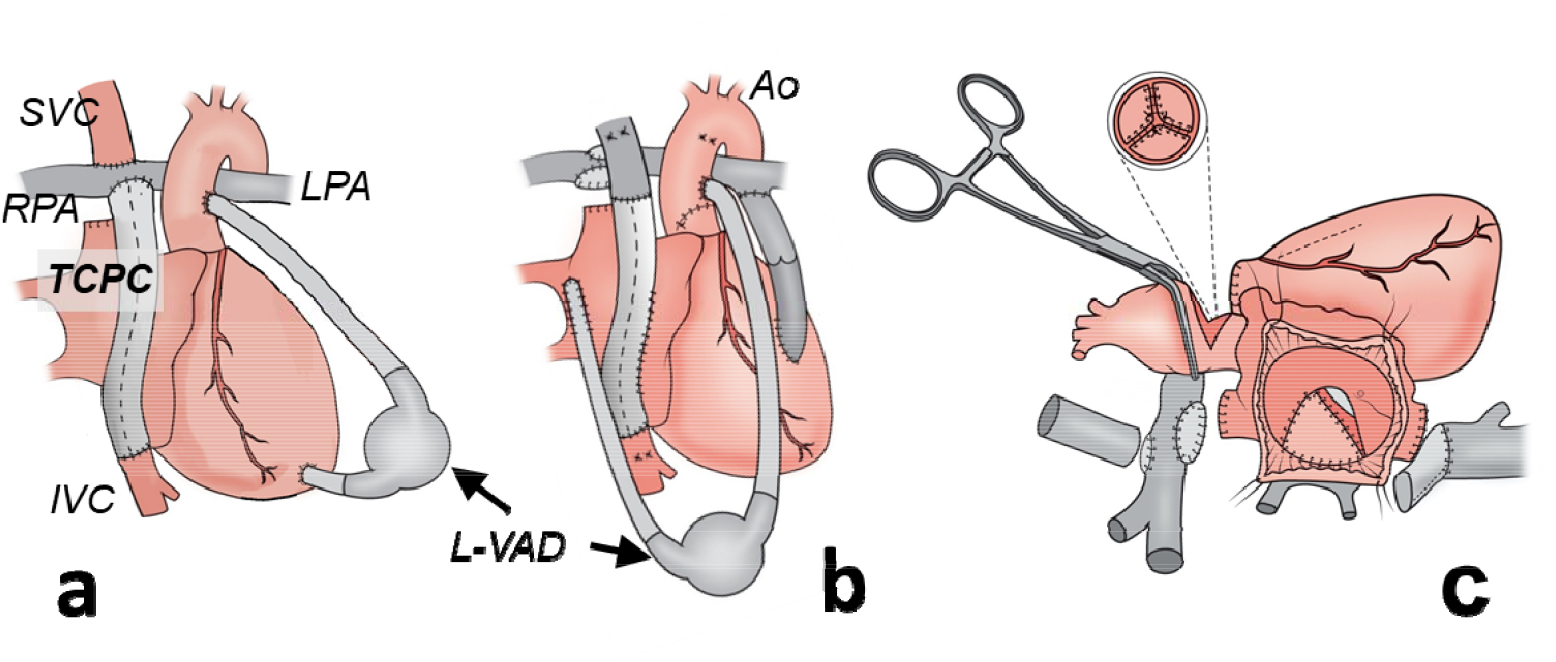
**(a)** Traditional left ventricle assist device (L-VAD) implantation configuration for systemic single-ventricle (SV) failure of the total cavopulmonary (TCPC) circulation. **(b)** Proposed configuration where the native SV is utilized as a pulsatile right heart. To achieve this Fontan (TCPC) tube graft is anastomosed to the right atrium, systemic circulation is maintained by L-VAD from pulmonary venous chamber to aorta (Ao), and pulmonary circulation is delegated to the SV via a conduit interposed between the SV and pulmonary artery bifurcation. **(c)** A sketch displaying the surgeons view is provided during the take-down of TCPC anastomoses, patch plasty of the pulmonary artery, creation of a posteriorly placed pulmonary venous chamber, and aortic valve closure through aortotomy. (LPA: Left pulmonary artery. RPA: Right pulmonary artery. IVC: Inferior vena cava, SVC: Superior vena cava)

To the best of our knowledge, there is no available technique in failing Fontan patients in whom the systemic circulation is maintained by a standard MCS, and the cavopulmonary circulation is delegated to the native SV as illustrated in Figure 1b. In this manuscript, analyses were performed for an optimal Fontan circulation, high PVR and ventricular dysfunction Fontan failures. These failure models were also examined under the conventional support and the proposed modification, using a clinical commercial HeartWare HVAD (Medtronic) device, *in vitro* and *in silico*. An established Fontan mock-up flow loop and a computational lumped parameter cardiovascular model (LPM) originally developed for Fontan circulation research were used to investigate a variety of clinically significant states.

## 2. Fontan Circulation States

### 2.1 Ideal Fontan Circulation (Fontan)

The ideal Fontan circulation state was modelled based on the Egbe et. al^23^ and our previous Fontan study^20^. SV was determined to have an ejection fraction (EF) of 44% and a stroke volume of 62 ml. Table 1 provides the characteristic clinical and anthropometric data based on the literature for the average age of an individual experiences Fontan failure. Cardiac parameters, systemic and pulmonary vascular resistances (SVR and PVR, respectively) were determined based on these typical represenative patient characteristics.

**Table 1.** Cardiovascular anthropometric data of an idealized Fontan patient. Compiled from literature and clinical experience^13,14^

| Variable | Data |
| --- | --- |
| Age, years | 15 |
| Height, cm | 150 |
| Body weight, kg | 50 |
| Body surface area, m<sup>2</sup> | 1.44 |
| Heart rate, bpm | 70 |
| Cardiac index, l/min/m<sup>2</sup> | 3 |
| Common atrial volume, ml | 70 |
| Systemic venous chamber volume, ml* | 45 |
| Pulmonary venous chamber volume, ml* | 25 |

\* indicates venous chamber volumes after partitioning of the common atrium (CA) employed in cases VD-switch, PVR-switch, bPVRtc-switch and bPVRtc-Pswitch.

For this baseline case, the cardiac index (CI) was set to 3.0 l/min/m^2^, and the systolic, diastolic, and mean aortic pressures were set to 100 mmHg, 67 mmHg, and 82 mmHg, respectively. Figure 2a and Figure 3a represent the LPM framework and *in vitro* Mock-up loop developed for this ideal (baseline) Fontan circulation, respectively.

**Figure 2.**
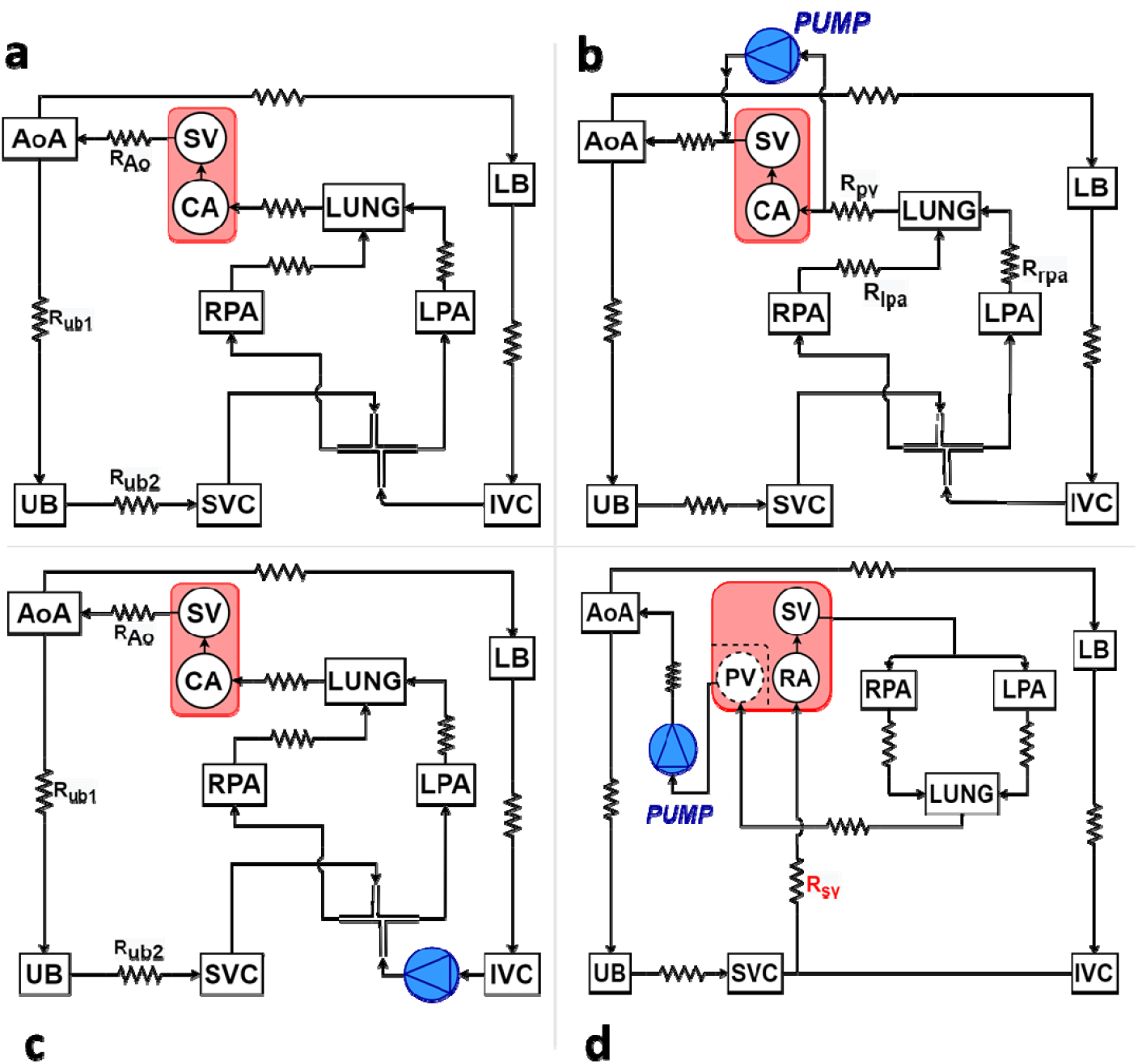
Electrical analog circulation networks analyzed in this study. **(a)** Healthy and failed Fontan circulations (Cases: *Fontan, VD* and *PVR*), **(b)** Conventional mechanical circulatory support (MCS) of ventricular dysfunction Fontan failure, Case: *VD-cmcs*. **(c)** Conventional MCS of increased PVR Fontan failure, Case: *PVR-cmcs* and **(d)** the proposed MCS modification cases *VD-switch* and *PVR-switch*. SV: single ventricle, CA: common atrium, AoA: aortic arch, LB (UB): lower (upper) body, IVC (SVC): inferior (superior) vena cava, RPA (LPA): right (left) pulmonary artery, LUNG: lungs, PV: posterior pulmonary venous (surgically separated from the CA with the dashed lines), RA: right atrium, R_Ao_: aorta resistance, R_lb_ (R_ub_): lower (upper) body resistances, R_TCPC_: TCPC resistance, R_rpa_ (R_lpa_): right (left) pulmonary artery resistance, R_sv_: systemic venous resistance, R_pv_: pulmonary venous resistance.

**Figure 3.**
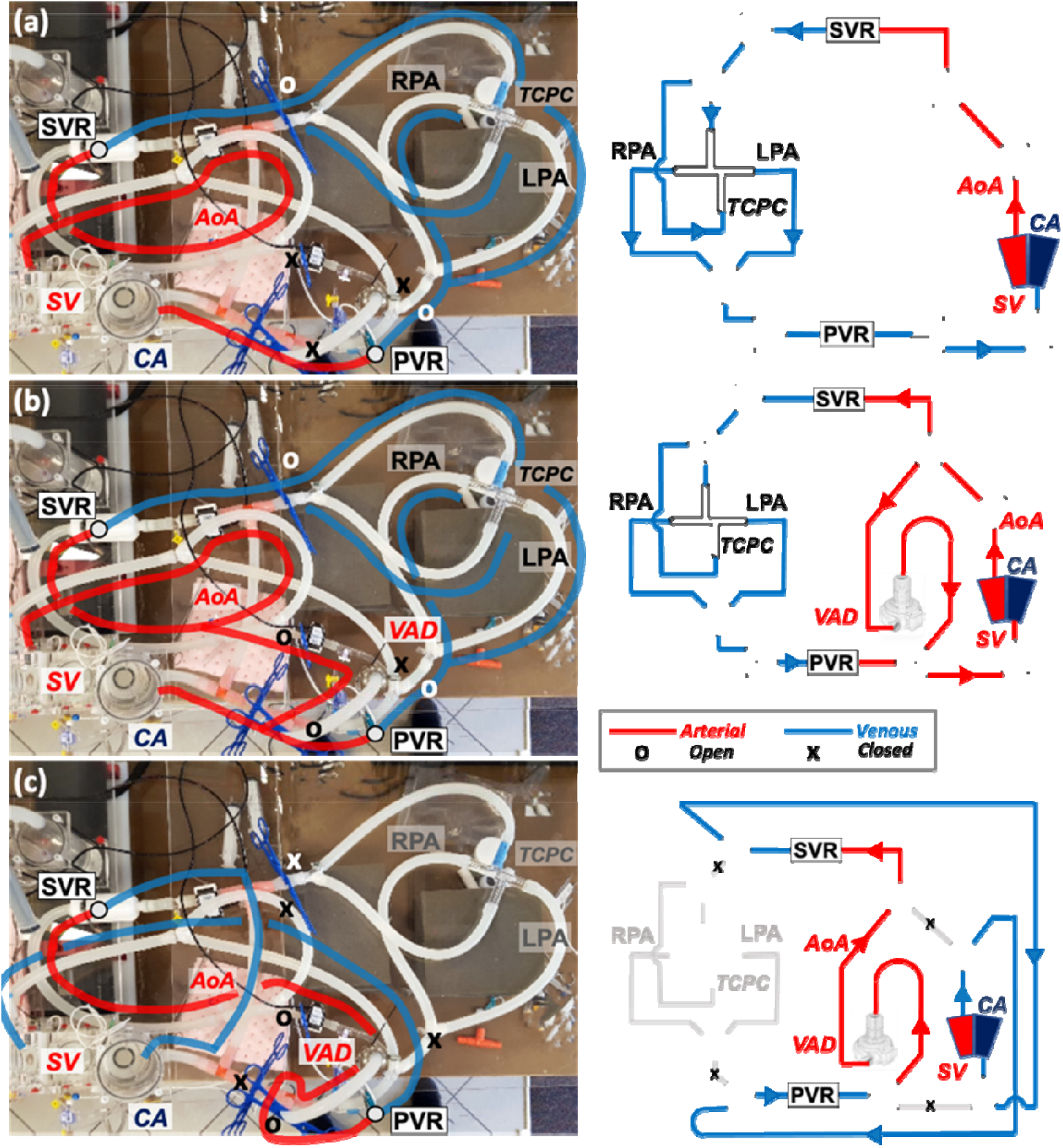
Fontan circulation cases replicated experimentally in our mock-up flow loop for *in-vitro* validation experiments. In a typical experimental run, cases are simulated in-sequence to allow direct comparison of measurements with each other. Starting with the **(a)** ideal, baseline *Fontan* circulation followed by the ventricular dysfunction failure case: *VD*, **(b)** MCS of ventricular dysfunction failure (*VD-cmcs*) and **(c)** the current modification where the MCS device operates as the left ventricle and the single-ventricle (SV) operates as the right ventricle (Case: *VD-switch*). *Red* and *blue* lines indicate the systemic and pulmonary circulations, respectively. *Gray* colored lines for the baseline Fontan network were removed in *VD-switch* configuration. SVR: Systemic vascular resistance, PVR: Pulmonary vascular resistance, RPA: Right pulmonary artery, LPA: Left pulmonary artery, Ao: Aorta, TCPC: Total cavopulmonary connection, CA: Common atrium, VAD: Ventricle assist device (HeartWare HVAD from Medtronic).

### 2.2 Fontan Failure, Ventricular Dysfunction (VD)

This case was generated based on moderate/severe ventricular dysfunction classification introduced by the New York Heart Association. Case *VD* was analyzed as two *VD-Ac* and *VD-Cr* cases considering the acute and chronical effects of the ventricular failure, respectively. In *VD-Ac*, stroke volume in *Fontan* was reduced from 62 ml to 38 ml, that leads to an EF of 26%. Therefore, sudden (acute) effect of the ventricular failure was simulated. In *VD-Cr*, physiologic response of cardiovascular system to the acute decrease in aortic pressure and CI was investigated. Such drop was compensated by increasing SVR index from 22.8 WU.m^-2^ to 27.6 WU.m^-2^.

### 2.3 Fontan Failure, Increased Pulmonary Vascular Resistance (PVR)

In *PVR* case, another common mode of Fontan failure was established through high PVR index based on Egbe et. al^23^. It was analyzed through cases *PVR-Ac* and *PVR-Cr* considering the acute and chronic effects of the high PVR index, respectively.

In *PVR-Ac*, PVR was increased from 1.65 WU.m^-2^ to 3.3 WU.m^-2^ to simulate the acute (sudden) effect of the PVR related Fontan failure. Altering the heart function or circulation parameters is not focused in this study in order to clearly isolate the sole effect of the proposed modification. Still an exercise case is provided to test off-design operation. Therefore, in *PVR-Cr*, as observed in Fontan patients, SVR index and systemic venous compliance were both increased by approximately 10% to replicate the physiological cardiovascular system response for preserving the cardiac output and systemic arterial pressure.

### 2.4 Conventional Mechanical Circulatory Support of the Ventricular Dysfunction Fontan Failure (VD-cmcs)

Case *VD-cmcs* corresponds to the conventional MCS (cmcs) of the Fontan failure state introduced in *VD-Cr*. In this configuration, MCS device supports the systemic circulation by working in parallel to the failing SV (EF=26%), as shown in Figure 2b and Figure 3b. The device was operated at 2560 rpm, which provided a flow rate of 2.82 l/min to support the ventricular dysfunction in terms of pressure and flow rate needs.

### 2.5 Conventional Mechanical Circulatory Support of the Fontan Failure with Increased Pulmonary Vascular Resistance (PVR-cmcs)

The conventional MCS support strategy intended for the high PVR Fontan failure model as introduced in *PVR-Cr (Section 2.3)* is simulated. Using our earlier cavopulmonary Fontan support framework^24^, MCS device was integrated between the systemic venous and pulmonary artery (PA), as shown in Figure 2c.

In *PVR-cmcs*, both continuous and pulsatile flow MCS device operation was investigated. For continuous flow support, MCS device was operated at a constant rotational speed of 2205 rpm. To impose the pulsatile operation condition, the rotational speed of MCS device was modulated sinusoidally (±400 rpm) during the operation. MCS device provided a flow rate of 3.0 l/min to decrease the central venous pressure (CVP) and support the cavopulmonary circulation in both operation conditions.

### 2.6 Proposed Modification, Tested for Ventricular Dysfunction Failure (VD-switch)

Case *VD-switch* represents the application of proposed modification to the Fontan failure mode introduced above as *VD-Cr (Section 2.2)*.

Here, a 25 ml volume was isolated surgically from the total common atrium (CA) volume (70 ml) to form a neo-pulmonary venous return chamber. The remaining portion of CA serves as a right atrium. Systemic venous return was redirected to the new right atrium, after detaching it from the conventional total cavopulmonary connection (TCPC). Thus, systemic and pulmonary circulations become parallel similar to a normal biventricular circulation. SV was connected to PA to maintain the pulmonary circulation. Systemic circulation was governed by the MCS device having an inlet draining from the neo-pulmonary venous chamber, yet its outlet was placed to the aorta, functioning like a native left ventricle. MCS device was set to work at 3300 rpm, which provided a total cardiac output of 4.95 l/min to the systemic circulation. In Figure 1b, a cartoon representation of the proposed modification is provided together with surgical details in Figure 1c. The corresponding circuit analogue is provided in Figure 2d as used in LPM computations.

Performance of the modification in case VD-*switch* was also investigated during the metabolic activity. To perform a simple leg activity (walking function), the exercise protocol introduced by Kung et. al.^25^ for Fontan patients was used. Based on this protocol, from rest (MET=0.65) to mild lower body exercise (MET=5), all parameters were remained constant except that the HR was increased from 66 bpm to 130 bpm and SVR was decreased by 15%.

### 2.7 Proposed Modification, Tested for the Increased Pulmonary Vascular Resistance (PVR-switch)

Here the proposed modification was applied to the high PVRI failure Fontan model introduced in *PVR-Cr (Section 2.3)*. MCS device governing the systemic circulation was operated at 3175 rpm corresponding to the flow rate of 4.32 l/min.

*PVR-switch* was also investigated for the activated compensatory mechanisms through *bPVRtc-switch*. In this case, heart rate was increased from 70 bpm to 120 bpm. Moreover, PA banding was applied to avoid an excessive increase in pulmonary pressure.

Analysis in *bPVRtc-switch* was repeated with the pulsatile MCS device operation introduced in *bPVRtc-Pswitch*. To observe the effect of pulsatility on the aortic flow and pressure, rotational speed of the device was modulated sinusoidally (±400 rpm) during the course of operation.

All these cases are summarized in Table 2.

**Table 2.** Clinically significant cases analyzed in this study using LPM and in vitro mock-up flow loop

|  |  |  |
| --- | --- | --- |
| **Fontan** |  | Ideal (optimally functioning) Fontan circulation. Corresponds to a time-point long after the 3<sup>rd</sup> stage surgery. i.e. immediate effects of establishing full single-ventricle (SV) circulation after surgery is recovered (eg. low post-op. cardiac output). |
| **VD** | **VD-Ac** | *Fontan Failure Mode 1:* SV systemic dysfunction associated failure of the Fontan circulation. Acute state is simulated, suddenly after the initiation of failure. |
|  | **VD-Cr** | *Fontan Failure Mode 1:* SV systemic dysfunction associated failure of Fontan circulation. Chronic state is simulated, long-term after the failure. |
| **PVR** | **PVR-Ac** | *Fontan Failure Mode 2:* High PVR associated failure of Fontan circulation. Acute state is simulated, suddenly after the initiation of failure. |
|  | **PVR-Cr** | *Fontan Failure Mode 2:* High PVR associated failure of Fontan circulation. Chronic state is simulated, long-term after the failure. |
| **VD-cmcs** |  | Conventional systemic support strategy for Fontan Failure Mode 1 (*VD-Cr*). Includes a clinical ventricle assist device (VAD) supporting systemic circulation. |
| **PVR-cmcs** |  | Conventional TCPC support strategy for Fontan Failure Mode 2 (*PVR-Cr*). Includes a Fontan assist device (FVAD) at the right-side, replacing TCPC conduit. |
| **VD-switch** |  | Proposed Fontan support strategy employing the native SV for Fontan Failure Mode 1 (*VD-Cr*). Hemodynamic performance is compared with Case *VD-cmcs*. |
| **PVR-switch** |  | Proposed Fontan support strategy employing the native SV for Fontan Failure Mode 2 (*PVR-Cr*). Hemodynamic performance is compared with Case *PVR-cmcs*. |
| **bPVRtc-switch** |  | Proposed support strategy for Fontan failure in (*PVR-Cr*) but with PA banding and tachycardia. |
| **bPVRtc-Pswitch** |  | Repetition of (*bPVRtc-switch*) with a pulsatile MCS device operation. Pulsatility is generated with sinusoidal rotational speed regulation. |

## 3. Methods

### 3.1 Proposed surgical modification

Through a traditional redo cardiac surgery approach, following aortic and selective bi-caval cannulation, cardiopulmonary bypass (CPB) is established. Under total CPB, the Fontan tube graft and superior cavopulmonary anastomosis are taken down from the right PA. The defects remained in the PA are patch repaired. The superior vena cava is anastomosed to the cephalic side of the Fontan tube graft. The aorta is cross clamped, and electromechanical quiescence is achieved by cardioplegia. Through right atriotomy, a patch is fashioned to fit for isolation of the pulmonary veins posterior to the left atrium. The aortic valve is closed through an aortotomy (Figure 1c). The Fontan tube graft is anastomosed side-by-side to the right atrium. Following a systemic ventriculotomy performed at the base of the heart, an appropriately-sized valved-conduit is interposed between the SV and the PA bifurcation. After priming of the MCS device, the outflow graft is anastomosed to the ascending aorta, and the inflow graft anastomosis to the left atrium is completed. Following removal of the aortic cross-clamp, the CPB flow is gradually reduced while the flow of the MCS device is increased synchronously (Figure 1c). By this way, left (via MCS device) and right sides (via SV) are totally separated like a biventricular circulation, providing a *Q*_*p*_/*Q*_*s*_ ratio of 1.

To investigate the proposed modification in a comparative manner, the same MCS device, HeartWare HVAD (Medtronic Inc, Fridley, Minnesota), was used for all *in silico* and *in vitro* cases. Therefore, pressure and flow hemodynamics are the main metrics to evaluate the performance of the modification subject to different circulation parameters or common disease states.

### 3.2 Lumped parameter Fontan circulation model

An established multi-compartmental LPM developed by our group for congenital heart disease research^26,27^ has been adopted to simulate the introduced Fontan circulation states. This model computes the pressure and flow patterns for key vascular components by representing them as compliance chambers and resistance vessels as given in Equation (1).

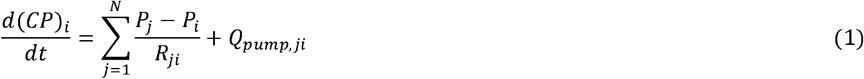

where *C* and *P* are the compliance and pressure of the compliant chamber represented by the index (*i* or *j*), respectively. *R* is the peripheral resistance of the vessel connecting the associated chambers. *N* is the number of lumped elements. *Q*_*pump*_ is the flow of the MCS device (Heartware HVAD) used in *in silico* and *in vitro* simulations at the time step (dt). Pump speed was determined based on the required *Q*_*pump*_, and remained constant during the analysis. Backward Euler method was used to iteratively solve the implicit formulation of Equation (1) using the fixed time step.

Compliance of the chambers are set as constant values based on the baseline patient profile (Table 1 and a file attached to the Data availability section). SV model function is modeled through the time-varying elastance concept introduced by Suga et. al.^28^ (*E*_*SV*_(*t*)) and the “double-Hill” function *E*_*n*_(*t*_*n*_) described by Stergiopulos et al.^29^ were used in Equation (2). Double-Hill function resolved ventricle characteristics reasonably well as presented in our earlier articles^27,30^.

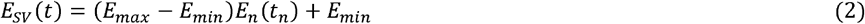

As for the patient’s anthropometric characteristics, all the simulations and measurements were based on a 15-year-old (1.44 m^2^) Fontan patient (Table 1), the age of which was considered as an approximate age of Fontan failure^20,23^. Cardiac functions, SVR and PVR were tuned according to the body size of the chosen patient profile.

### 3.3 In-Vitro Mock-Up Circulation

The LPM simulations were validated against our pediatric pulsatile mock-up Fontan flow loop for the ideal/functional (*Fontan*), ventricular dysfunction (*VD*), conventionally assisted ventricular dysfunction (*VD-cmcs*) and the proposed modification assisted ventricular dysfunction (*VD-switch*) Fontan failure circulations. This bench-top circulation system included a compliant ventricular phantom and computer-controlled pulse-duplicator (SuperDup’r, Vivitro Systems Inc, BC, Canada), which set the pulsatile flow rate by adjusting stroke and stroke volume, as described in our previous publications^20,31^. As per our experimental protocol, Fontan circuits were generated in sequence using clamps and Y-branches without significantly altering the circuit parameters or stopping the piston-pump, as seen in Figure 3. Thus, all the tested cases were comparable with each other and corresponded to the early acute changes of the proposed configuration. *In vitro* compartment parameters were adjusted representing the combination of lumped chambers in Figure 2 for sake of simplicity, so that these lumped chambers are not separately represented in Figure 3.

Our previously used standard 13.3 mm diameter one-degree offset based on the chest MRI of a Fontan patient TCPC connection made of glass was attached to inferior/superior vena cava and right/left PA compliance chambers^32^. Two clamp-on ultrasonic pulsatile flow transducers, namely a 3PXL to the pump outlet and an 8PXL to the aorta were then connected to TS410 flow modules (Transonic Systems Inc., Ithaca, New York), which were placed downstream of the SVR and MCS device outlets. Pressure measurements (Deltran 6200, Utah Medical Products Inc., Midvale, Utah) were obtained from the SV, aorta, MCS device outlet, pulmonary venous chamber and systemic venous bed. CVP was obtained from the pressure sensor placed on the inferior vena cava part of the Y-branch right after SVR. Measurements were recorded using the Lab-chart data acquisition unit (AD Instruments, Colorado Springs, Colo). Distilled water was used in the *in vitro* experiments at room temperature.

## 4. Results

### 4.1 In vitro validation of lumped parameter circulation model

Ideal and ventricular dysfunction Fontan failure models (*Fontan, VD, VD-cmcs* and *VD-switch*) were replicated exactly in an experimental mock-up circulation flow loop for numerical validation. The pressure drop based on the inertance of the tubes representing the cardiovascular elements in our *in vitro* setup was also calculated. The maximum inertance was observed at the tube representing the aorta, as expected, which has a length of 0.03 m and a radius of 0.007 m. The inertance of this section results a maximum pressure drop of only 4% of the mean aortic pressure. Likewise, the inertance based on pressure drop in the venous tubing components are observed to be 10^−3^ mmHg, primarily due to the low pulsatility. According to these observations, in silico and in vitro results were compared and agreed even though neglecting the inertance effect for simplicity.

For all these cases, measured aortic pressure waveforms demonstrated acceptable agreement with *in silico* LPMs computations, as shown in Figure 4. There is a small phase difference observed between the waveforms, which is based on the HR difference of 4% between *in silico* and *in vitro* simulations.

**Figure 4.**
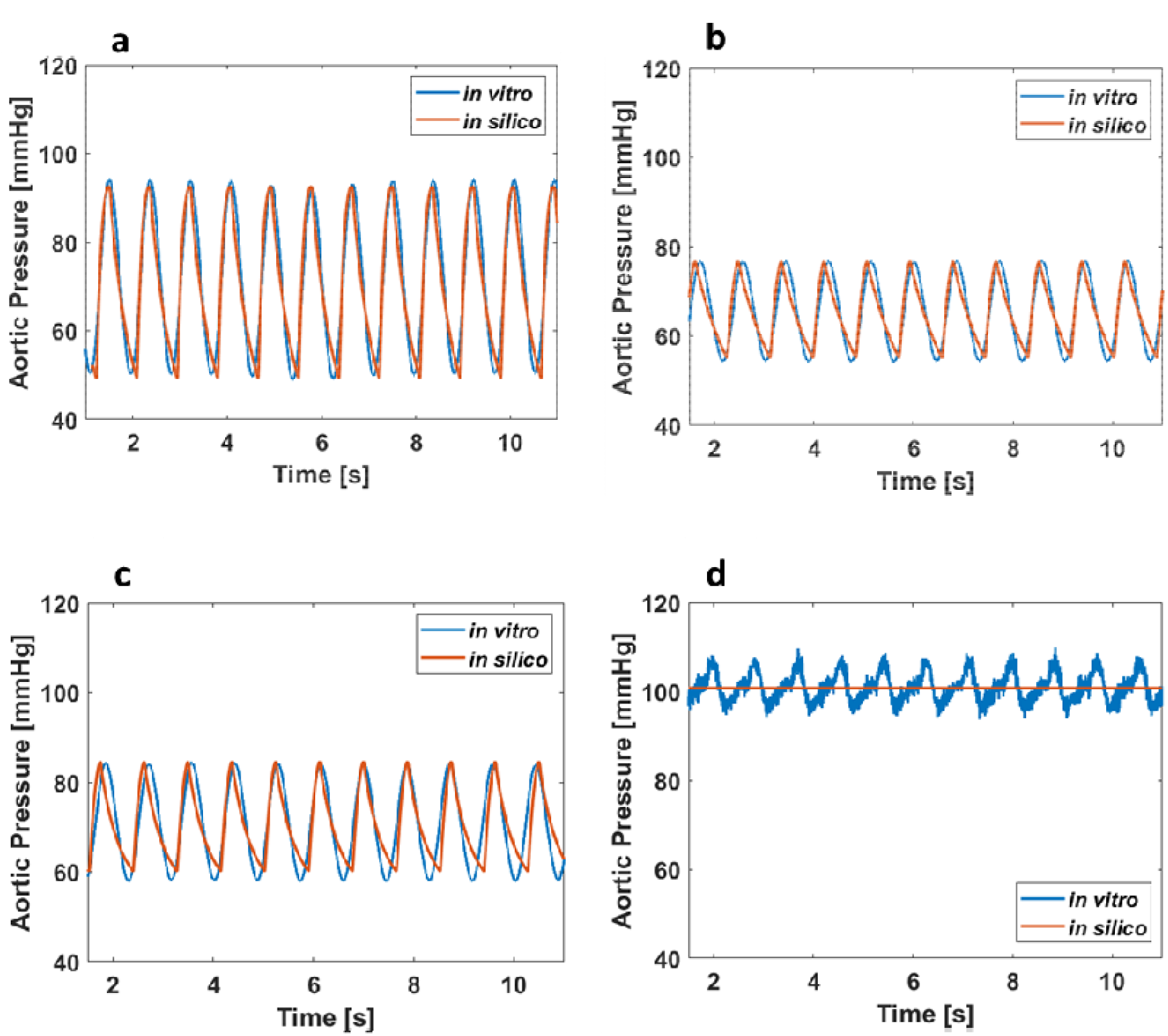
Representative hemodynamic waveforms for experimental validation. Aortic pressure waveforms obtained *in vitro* (mock-up flow loop measurements) and *in silico* LPM model are plotted for the circulation cases of **(a)** the healthy *baseline* (for ventricular dysfunction cases); *Fontan*, **(b)** ventricular dysfunction; *VD*, **(c)** conventional mechanical circulatory support of VD; *VD-cmcs* and **(d)** when the right-ventricle is used a right-heart in VD as proposed in this manuscript; *VD-switch*. All waveforms presented are recorded after the mock-up flow loop is stabilized and operating at the steady pulsatile hemodynamics.

Likewise, simulated hemodynamics were also validated through pulsatile *in vitro* measurements. For the SV pressure, simulated and measured values in *Fontan* and *VD-switch* matched almost exactly. On the other hand, almost 6% difference between *in silico* and *in vitro* measurements is achieved both in *VD* and *VD-cmcs*. In terms of CVP, *in silico* results agreed well with *in vitro* measurements for *VD* and *VD-switch*. However, CVP revealed a difference of 11% and 17% between the *in silico* and *in vitro* for *Fontan* and *VD-cmcs*, respectively. In *VD-switch*, MCS device pressure was recorded to be 101 mmHg in both *in silico* and *in vitro* analyses, yet it revealed a difference of 3% in CI values. In *VD-cmcs*, the CI of 3.43 l/min/m^2^ was observed in both *in silico* and *in vitro* simulations. This discrepancy (max. 6.5% observed between *in silico* and *in vitro*) is due to the MCS device pressure adjusted to provide the same CI. There was no significant difference (<1 mmHg) in pulmonary venous pressures between the *in vitro* and *in silico* models in all cases.

In all cases, a continuous flow MCS device, Heartware HVAD (Medtronic) was used. However, a pulsatile aortic pressure waveform was observed through *in vitro* analysis in *VD-switch*, as seen in Figure 4d. In this case, the pressure sensor placed at MCS device output in *in vitro* experiments was affected by the pulsating piston ventricle of the mock-up loop (Figure 3c), which is the reason of such pulsatile aortic waveform. Therefore, even though the same mean aortic pressure was achieved, different waveforms was observed in *in silico* and *in vitro* results.

### 4.2 Fontan Failure-1, Ventricular Dysfunction Model

Results associated with ventricular dysfunction models (*VD, VD-cmcs* and *VD-switch)* are shown in Table 3.

**Table 3.** Simulated hemodynamic parameters of the ventricular dysfunction Fontan failure state.

|  | (VD-Cr)<br>Ventricular Dysfunction<br>Fontan Failure<br>(Chronic) | (VD-cmcs)<br>Conventional Systemic MCS,<br>Ventricular Dysfunction Fontan<br>Failure | (VD-switch)<br>Proposed Modification,<br>Ventricular Dysfunction Fontan<br>Failure |
| --- | --- | --- | --- |
| HR [bpm] | 66 | 66 | 66 |
| EF [%] | 26 | 26 | 26 |
| Stroke volume index [ml/m<sup>2</sup>] | 26 | 22 | 43 |
| CO [l/min] | 2.47 | 4.93* | 4.95 |
| CI [l/min/m<sup>2</sup>] | 1.72 | 3.42 | 3.44 |
| MCS device output [l/min] | n/a | 2.82 | 4.95 |
| SBP [mmHg] | 77 | 85 | 101 |
| DBP [mmHg] | 55 | 60 | 101 |
| MBP [mmHg] | 65.5 | 71 | 101 |
| CVP [mmHg] | 10.8 | 11.35 | 12 |
| PAP [mmHg] | 9 | 7.85 | 17.5 |

Abbreviations: CI: cardiac index, CMRI: cardiac magnetic resonance imaging, CO: cardiac output, CVP: central venous pressure, DBP: diastolic blood pressure, EF: ejection fraction, HR: heart rate, PAP: pulmonary artery pressure, SBP: systolic blood pressure, MBP: mean blood pressure, MCS: mechanical circulatory support. Note that the ideal Fontan circulation hemodynamic parameters to impose ventricular dysfunction are nearly 10% higher than the used one for increased PVR model. Thus, this condition is not additionally given in this table. Figure 4a represents the waveform obtained for this *baseline* condition. \*CO represents the total blood flow provided by the MCS device and SV.

In *VD*, EF of 26% led to a decrease in CI and aortic pressure by 1.7 l/min/m^2^ and 5 mmHg, respectively. Since pumping energy (stroke volume) of the SV was reduced to decrease EF, CVP also decreased by 4 mmHg. Correspondingly, PA pressure decreased slightly from 11 mmHg to 9 mmHg.

In *VD-cmcs*, CI was increased by 1.7 l/min/m^2^ with the conventional systemic support of MCS device. It augmented both the aortic and ventricular pressures by 5 mmHg and 4 mmHg, respectively. CVP pressure was barely increased due to the implantation configuration of MCS device in this case (Figure 3b), unlike the conventional cavopulmonary support.

In *VD-switch*, MCS device directly reflects the pulmonary flow to systemic circulation. In pulmonary side, although the stroke volume of SV was nearly doubled, EF of it remained constant. Mean SV pressure was simulated as 18.5 mmHg. Additionally, PA pressure increased from 10.7 mmHg to 17.5 mmHg (39%). CVP was decreased by 2 mmHg (13%). *In silico* waveforms simulated for the ventricular dysfunction model are shown in Figure 5. As expected, suction effect of MCS device caused a decrease in pulmonary venous pressure to −1.2 mmHg, almost collapsing the *in-vitro* compliance chamber as seen in Figure 5a. Nevertheless, CI and aortic pressure increased by 1.7 l/min/m^2^ (doubled) and 25 mmHg (38%), respectively, to compensate for the low EF.

**Figure 5.**
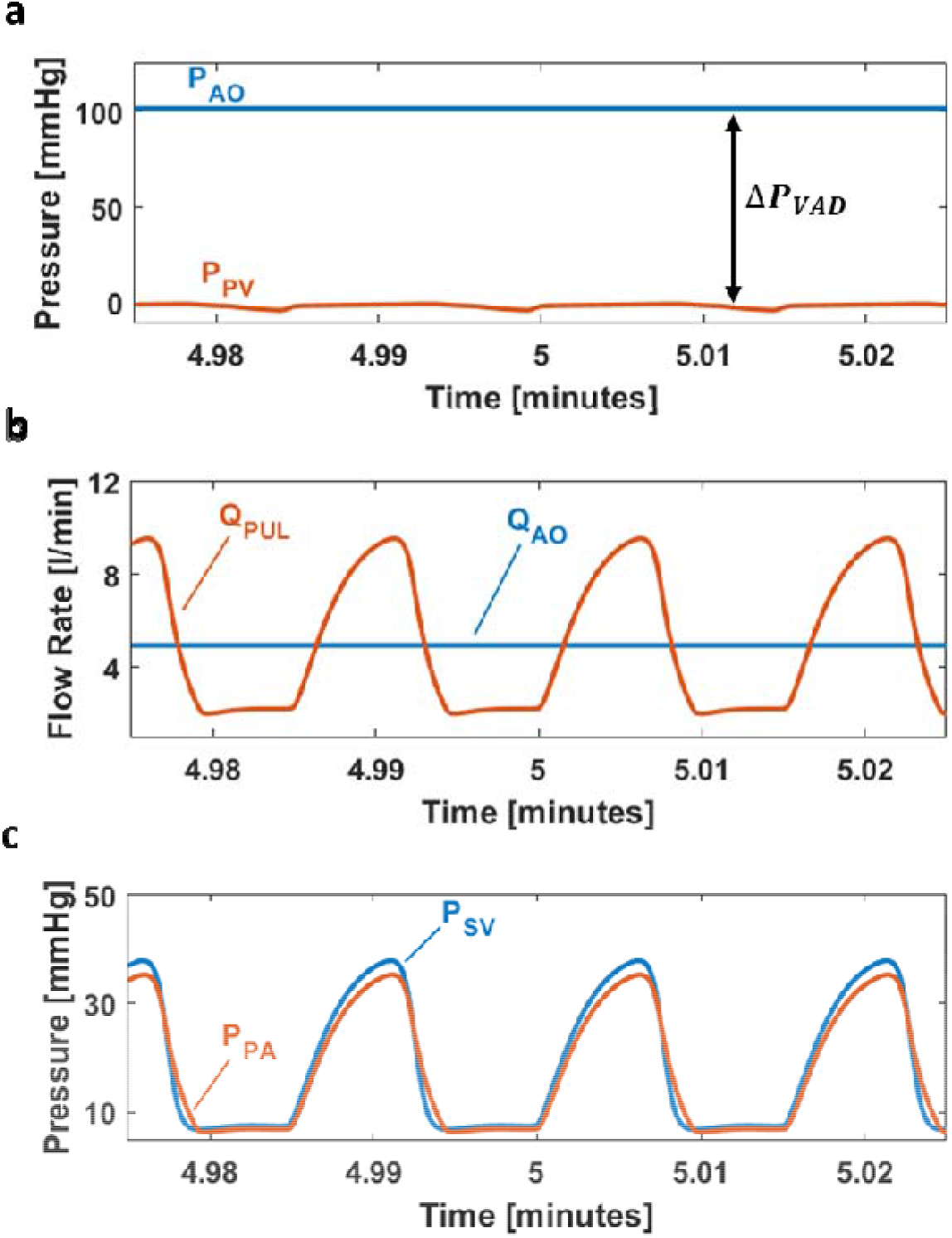
Hemodynamic waveforms computed when the single-ventricle (SV) is spared as a right-heart during the MCS of ventricular dysfunction Fontan failure mode (Failure Mode 1). **(a)** Aortic (P_AO_) and pulmonary venous (P_PV_) chamber pressures, **(b)** Aortic (Q_AO_) and pulmonary (Q_PUL_) flow waveforms with the MCS **(c)** Single ventricular (P_SV_) and pulmonary artery (P_PA_) pressure waveforms. All waveforms presented are recorded after the mock-up flow loop is stabilized and operating at the steady pulsatile hemodynamics.

### 4.3 Fontan Failure-2, Increased Pulmonary Vascular Resistance Model

Hemodynamics associated with the high PVR index Fontan failure cases (*PVR-Ac/Cr, PVR-cmcs, PVR-switch, bPVRtc-switch* and *bPVRtc-Pswitch*) are presented in Table 4.

**Table 4.** Simulated hemodynamic parameters of the increased PVR Fontan failure state.

|  | *(Fontan)*<br>Ideal Fontan<br>Circulation | *(Fontan)*<br>Ideal Fontan<br>Circulation<br>(Egbe et al.[20]) | *(PVR-Ac)*<br>Increased<br>PVR Fontan<br>Failure<br>(Acute) | *(PVR-Cr)*<br>Increased<br>PVR Fontan<br>Failure<br>(Chronic) | *(PVR-cmcs)*<br>Conventional<br>MCS,<br>Increased<br>PVR Fontan<br>Failure | *(PVR-switch)*<br>Proposed<br>Modification,<br>Increased PVR<br>Fontan Failure | *(bPVRtc-switch)*<br>Proposed<br>Modification<br>(Tachycardia and<br>pulmonary banding) | *(bPVRtc-Pswitch)*<br>Proposed<br>Modification<br>(Pulsatile operation,<br>tachycardia and<br>pulmonary banding) |
| --- | --- | --- | --- | --- | --- | --- | --- | --- |
| HR [bpm] | 70 | 69 ± 7 | 70 | 70 | 70 | 70 | 120 | 120 |
| EF [%] | 44 | Echo - 43 ± 4<br>CMRI - 47 ± 6 | 42 | 42 | 42 | 43 | 27 | 27 |
| Stroke<br>volume index<br>[ml/m<sup>2</sup>] | 43 | Echo - 46 ± 6<br>CMRI - 45 ± 6 | 37 | 40 | 45 | 43 | 25 | 27 |
| CO [l/min] | 4.32 | 5.76 (4.15 - 7) | 3.69 | 4.05 | 4.55 | 4.32 | 4.32 | 4.36 |
| CI [l/min/m<sup>2</sup>] | 3 | 3.2 (2.3 - 3.9) | 2.56 | 2.81 | 3.16 | 3 | 3 | 3.02 |
| MCS device<br>output [l/min] | n/a | - | n/a | n/a | 3 | 4.32 | 4.32 | 4.36 |
| SBP [mmHg] | 100 | - | 90 | 100 | 107 | - | - | 81 |
| DBP [mmHg] | 67 | n/a | 62 | 65 | 69 | - | - | 77 |
| MBP [mmHg] | 82 | 81 ± 5 | 75 | 82 | 88 | 78 | 78 | 79 |
| CVP [mmHg] | 14 | 15 ± 4 | 16.25 | 18 | 15 | 8.7 | 9.1 | 9.25 |
| PAP [mmHg] | 10 | 10 ± 2 | 12.15 | 13.5 | 15 | 42 | 37 | 35 |

Abbreviations: CI: cardiac index, CMRI: cardiac magnetic resonance imaging, CO: cardiac output, CVP: central venous pressure, DBP: diastolic blood pressure, EF: ejection fraction, HR: heart rate, PAP: pulmonary artery pressure, SBP: systolic blood pressure, MBP: mean blood pressure, MCS: mechanical circulatory support.

In *PVR-Ac*, a decrease in CI by 0.35 l/min/m^2^ and mean aortic pressure by 7 mmHg was observed. CVP increased from 14 mmHg to 16.25 mmHg at this acute stage of failure. Elevated CVP also increased the PA pressure by 3 mmHg. In *PVR-Cr* representing the chronic condition, hemodynamic mechanisms tended to pull the systemic parameters to the ideal Fontan circulation by increasing the SVR index. Thus, CVP elevated from 14 mmHg to 18 mmHg, which led to the serious failure of Fontan circulation in time.

In *PVR-cmcs*, CVP decreased by 3 mmHg (16.7%) with the cavopulmonary conventional MCS. Therefore, CI increased by 0.35 l/min/m^2^ (12.5%) and the mean aortic pressure increased by 6 mmHg (7.3%). Using the MCS device as a right ventricle in this case also increased the PA pressure from 13.5 mmHg to 15 mmHg (11.1%).

In *PVR-switch*, the mean aortic pressure was observed as 78 mmHg with the CI of 3.0 l/min/m^2^. The EF of the SV, which governs the pulmonary circulation remained as 43% while the MCSD propelled 4.32 l/min of blood to the systemic circulation. As a result, the desired systemic hemodynamic measurements as per aortic pressure and CI were obtained, which led to a decrease in CVP to 8.7 mmHg (38%). However, the PA pressure was excessively elevated to 42 mmHg as expected.

In *bPVRtc-switch*, such pulmonary hypertension was aimed to ease through tachycardia and PA banding, which led to a decrease in EF from 43% to 27%. Correspondingly, the PA pressure decreased to 37 mmHg from 42 mmHg, while the aortic pressure and CI remained nearly constant as in *PVR-switch*. Additionally, the CVP slightly increased to 9.1 mmHg.

In *bPVRtc-Pswitch*, an approximately ±3 mmHg of pulsatility was generated in the pulmonary flow through replacement of the continuous flow MCS device with a pulsatile one. Since the stiffness of the aortic compliance chamber is less than the PA chamber in the current LPM, only ±2 mmHg of aortic pressure pulsatility was observed. The pulsatility in pulmonary flow was observed as ±1.15 l/min. Although the application of a pulsatile MCS device did not significantly affect the mean aortic and PA pressures, it just provided a more physiological systemic flow waveform. Figure 6 shows the effect of pulsatile MCS device on the hemodynamic waveforms in *bPVRtc-Pswitch*. Figure 6a represents the aortic and pulmonary venous pressure waveforms under the pulsatile pump effect. Assisted aortic and pulmonary flow waveforms are shown in Figure 6b. Figure 6c demonstrated that the pulsatility generated by MCS device varied the blood flow rather than the compliance chamber pressure. Pulsatility of the PA pressure was approximately ±9 mmHg. However, tachycardia and PA banding nearly eliminated the reverberation of it as seen in Figure 6d.

**Figure 6.**
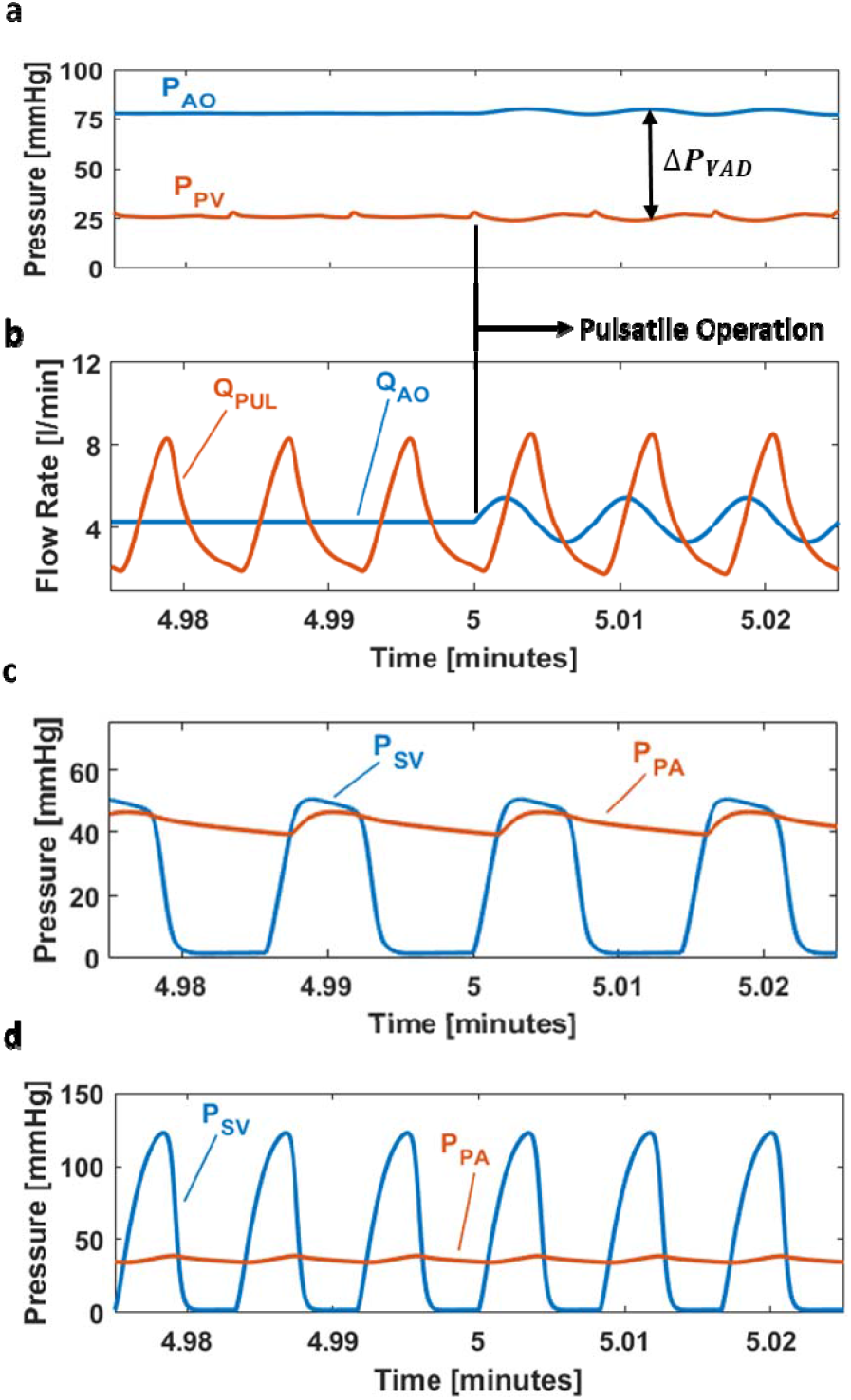
Hemodynamic waveforms computed when the single-ventricle (SV) is spared as proposed during the MCS of high PVR associated failure of Fontan circulation (Failure Mode 2). **(a)** Aortic (P_AO_) and pulmonary venous (P_PV_) chamber pressures, **(b)** Aortic (Q_AO_) and pulmonary (Q_PUL_) flow waveforms with the pulsatile MCS device operation, banding and tachycardia (Case *bPVRtc-Pswitch*), **(c)** Single ventricular (P_SV_) and pulmonary artery (P_PA_) pressure waveforms without banding and tachycardia (Case *PVR-switch*), **(d)** Single ventricle (P_SV_) and pulmonary artery (P_PA_) pressure waveforms with banding and tachycardia (Case *bPVRtc-switch*). Rotational speed modulation (±400 rpm) to produce pulsatility starts at time= 5 minutes. All waveforms presented are recorded after the mock-up flow loop is stabilized and operating at the steady pulsatile hemodynamics.

## 5. Discussion

The literature includes a variety of MCS device implantation concepts for failing Fontan circulation^15–19,33,34^. The micro-axial and vicious impeller flow devices provide mechanical augmentation of the TCPC^17–19^ with low blood damage. Although these systems are effective in reducing CVP^17^, if they fail or shut down they obstruct venous flow^18^. Most importantly, the chronic long-standing privation of the pulsatile pulmonary blood flow in Fontan circulation is considered the main reason of the Fontan failure through altered endothelial-dependent vasorelaxation response and depressed expression of endothelial nitric oxide synthase^18,24^. Use of a right-sided continuous flow MCS device continues this vicious cycle albeit at reduced venous pressure levels. The proposed concept has potential to provide a pulmonary antegrad flow reaching physiological pulsatility levels, which would also decrease pulmonary capillary recruitment and pulmonary vascular impedance^35^, eliminating the detrimental effects of non-pulsatile pulmonary flow. This would presumably improve the condition of the patient during bridge-to-heart transplantation.

Figure 5b and 5c show that the proposed modification for the ventricular dysfunction Fontan failure (*VD-switch*) provided a mean PA pressure and pulmonary waveform very similar to the native pulmonary pressure and flow, respectively. Additionally, simulated mean pressure levels of the SV used at the pulmonary side was observed to be very similar to the native right ventricle. Therefore, the proposed modification might be a potential solution providing a more physiological pulmonary flow than the conventional Fontan failure MCS. Moreover, subject to the significantly lower afterloads of the right-side, the failing SV at the pulmonary position functions at a more desirable operating point, even though it is insufficient to address the higher systemic circulation loading. During a mild lower body exercise, as simulated by our LPM model, Case *VD-switch* provided the required increase in CO and PA pressure by 28% and 35%, respectively. MCS device output and aortic pressure barely increased (maximum ∼5%). Therefore, it was observed that the HR increased during metabolic activity affects the PA pressure more than the aortic pressure. This is expected since SV governs the pulmonary circulation while LVAD characteristics governs the systemic circulation more than the SV function.

*PVR-Cr* represents a chronic increased PVR Fontan failure state, in which the CVP is maintained at 18 mmHg, which is far over the optimal limit (14 mmHg) for an ideal Fontan patient^36^. Based on the literature, even a 2 to 6 mmHg of support is effective in assisting cavopulmonary circulation and reducing CVP that will delay Fontan failure^18,19^. Correspondingly, a 3 mmHg drop in CVP achieved through conventional cavopulmonary support simulations (*PVR-cmcs*) agrees with the literature. Furthermore, the proposed modification cases with increased PVR Fontan failure (*PVR-switch* and *bPVRtc-Pswitch*) provided further decrease in the CVP (by 9.3 mmHg and 8.75 mmHg, respectively), which indicates the effectiveness of this concept.

In the modified biventricular assistance of Nathan et al.^33^, both the superior and inferior cavopulmonary anastomoses are disconnected from the PA, and a new right-sided venous reservoir is created for the systemic venous inflow. Although device thrombosis was the main reason of their patient’s death, 29% mortality rate was reported by de Rita et. al.^34^. Although the concept of creating a biventricular physiology by separating the systemic and pulmonary venous return resembles our proposed modification, the major difference of our concept is that a biventricular support is reconstructed by using one left-sided MCS device. By this way, the complexity pertaining to the use of two separate devices is avoided, which, in our opinion, is another advantage of the current concept.

Woods et al.^16^ introduced the use of posterior pexy of the tricuspid sub-valvar apparatus to maintain an adequate inflow from the SV in right-heart morphology in addition to the ventricular cannulation for inflow^14–16,33,34^. Thus, the presumed advantage of the current modification is avoidance of inflow ventricular cannulation. However, the current concept comprises a ventriculotomy for ventricle-to-PA conduit interposition, which arouses concern regarding whether the ventriculotomy-applied ventricle could maintain the pulmonary circulation or not in patients, who already had mildly reduced EF and high PVR (*PVR-Ac* and *PVR-Cr*). Pertinently, it is hypothesized that this mildly-failed SV was already supporting the whole circulation before transitioning and would be adequate for the right side. This conclusion was also supported by the findings of the proposed modification for increased PVR model (*PVR-switch*). In this case, SV redundantly supported the pulmonary circulation and moreover, led to a considerable increase in PA pressure reaching to 42 mmHg since the pumping capacity of the even mildly failed SV was considerably higher than a typical right ventricle subject to the Frank-Starling mechanism. The rise in PA pressure is one of the existing clinical challenges of patients who received a left-ventricular, or bi-ventricular assist device support^37^. In order to overcome the excessive rise in PA pressure, *bPVRtc-switch* was created featuring a banded or an undersized RV-to-PA conduit. This increased resistance led to a one third decrease in PA pressure from 42 to 37 mmHg. On the other hand, in *VD-switch* when a severely failed SV is used in the pulmonary position, excessive pulmonary pressures was not observed due to the lower pumping capacity. This case also provided a significant pulmonary flow pulsatility compared to the increased PVR failure support with the proposed modification, as summarized in Figures 5c and 6d. Therefore, a failed SV utilized in the pulmonary circulation, which is common for the target clinical problem, will yield healthy hemodynamics at the right-side compared to a mildly reduced SV function in high PVR case. Accordingly, results revelated that the hemodynamic benefit is more pronounced in patients with low EF, enabling clinicians to utilize even a severely failing ventricle in the systemic circulation on the pulmonary side.

The literature comprises applications in which the inflow is directly taken from the left atrium^34^. As a surgical strategy, in case of an inadequate inflow from the apical SV, de Rita et al.^34^ switched the inflow cannula to the common atrium. In respect thereof, the main foreseen constraint of our modification is the capacity of the pulmonary venous chamber, which was determined to be 25 ml. Even a slight obstruction to the pulmonary venous chamber would lead to device malfunction and death. Thus, the creation of the pulmonary venous chamber was thought to be the crux of the current modification. Even so, the suctioning effect of a continuous flow MCS device would reflect to the pulmonary venous bed and inevitably lead to collapse. Thus, it can clearly be predicted that the current concept is not suitable for patients with stage III-palliated hypoplastic left heart syndrome due to their innately small left atrial chamber. Additionally, presence of a Damus-Kay-Stansel root would preclude the creation of an SV-to-PA conduit interposition.

The most important constraint of the current concept is fail to reach the anticipated levels of systemic circulatory support due to ineffective left atrial unloading. As a back-up plan for this malady, creating a systemic venous compartment within the Fontan tube graft through taking-down the Fontan graft-right atrial anastomosis, and switching the inflow cannula to the Fontan graft, creating a pulmonary venous atrium through removing the intra-atrial patch, connecting the outflow cannula to the PA or the conduit, and re-opening the aortic valve can be applied. By this way, biventricular circulation can be maintained but with an assisted pulmonary circulation. Another foreseen concern of the current modification is the complexity of the surgical procedure with potentially long cardiopulmonary by-pass times in a patient with an already impaired ventricular function. The concept in creating a left atrial chamber is similar to the already existing Senning procedure^38^, which supports the clinical viability of proposed modification despite its complexity.

The surgical complexity necessitates another back-up plan if the proposed modification fails. In case of SV failure, rather than using another RVAD replacing the SV, which evolves the system to a BiVAD circulation, the modification will be reversed to a Fontan circulation with TCPC and an LVAD.

Similar to continuous flow MCS devices^15^, the Berlin Heart Excor can be applied to a wide range of patients from infants to adults. A pulsatile extra-corporeal MCS device seems to be more convenient for the current modification due to the effective decompression during pump diastole that is only 60% of each pump cycle^15^. By this way, the continuous suctioning effect of a continuous flow device could be avoided. In addition, the average 40% of blood left in the relatively small pulmonary venous chamber can further prevent its collapse.

The LPM and mock-up flow loop indicated that the proposed arrangement would work and satisfy the biventricular pressure and flow levels. One caution is the requirement of a relatively low venous compliance level. Otherwise, the suction generated by the MCS device would lead to collapse in pulmonary venous chamber. Therefore, a high systemic (MCS device) pressure reaching 110 mmHg to 120 mmHg might be challenging to keep this chamber open in the proposed modification. Although we observed the maximum pressure drop in the pulmonary venous chamber as almost −1 mmHg, the clinically-recorded limit of negative pressure in vacuum-assisted venous return is between −20 and −40 mmHg^39^. Therefore, it is predicted that the collapse risk is minimal during the clinical application of proposed modification.

However, an average systemic pressure around 90 mmHg to 95 mmHg can easily be achieved with no pulmonary venous suctioning effect.

## 6. Conclusion

Achieving an optimal management strategy for failing Fontan circulation by MCS devices is an ongoing effort. A consensus on the ideal MCS device implantation strategy has not yet defined due to the limited data and clinical experience as well as large patient-to-patient variation. Even though the detailed in vitro and in silico simulations of the proposed fictional concept with the use of an actual continuous flow device showed encouraging results, our modification is originally planned for the long-term support as a bridge to transplantation. Therefore, it is essential to test this proof-of-concept idea in an animal model. Thus, specially targeting the low resource settings with limited access to MCS devices and heart transplantation, sparing the native ventricle as a right-heart support will provide a novel perspective for MCSD device implantation in failed Fontan patients.

## Data availability

There are no restrictions on the availability of materials or information. The datasets generated and/or analyzed during the current study are available via *https://doi.org/10.5281/zenodo.6300829*. For any questions, please contact corresponding authors.

## Acknowledgements

We would like to express our gratitude to Medtronic for providing a loaner HeartWare MCS pump during the *in vitro* experiments and Elsevier Illustration Service for the drawing of Figure 1.

## Author contributions

ES, ONT, YA, MO, KP hypothesized and introduced the proposed concept. CY, BA, KP designed and conducted computational and experimental work. All authors wrote and edited the manuscript text.

